# Interaction of the mechanosensitive microswimmer *Paramecium* with obstacles

**DOI:** 10.1101/2022.12.20.521225

**Authors:** Nicolas Escoubet, Romain Brette, Lea-Laetitia Pontani, Alexis M. Prevost

## Abstract

In this work, we report investigations of the swimming behavior of *Paramecium tetraurelia*, a unicellular microorganism, in micro-engineered pools that are decorated with thousands of cylindrical pillars. Two types of contact interactions are measured, either passive scattering of *Paramecium* along the obstacle or avoiding reactions, characterized by an initial backward swimming upon contact, followed by a reorientation before resuming forward motion. We find that avoiding reactions are only mechanically triggered about 10% of the time. In addition, we observe that only a third of all avoiding reactions triggered by contact are instantaneous while two thirds are delayed by about 150 ms. These measurements are consistent with a simple electrophysiological model of mechanotransduction composed of a strong transient current followed by a persistent one upon prolonged contact. This is in apparent contrast with previous electrophysiological measurements where immobilized cells were stimulated with thin probes, which showed instantaneous behavioral responses and no persistent current. Our findings highlight the importance of ecologically relevant approaches to unravel the motility of mechanosensitive microorganisms in complex environments.

## I. INTRODUCTION

In their natural habitat, microorganisms evolve in complex environments that are characterized by the presence of obstacles with different shapes, sizes and mechanical properties. Therefore, foraging motile microorganisms have to swim efficiently in response to various stimuli such as chemical or light gradients. In this context, ciliated microorganisms have the ability to sense the properties of their environment through mechanosensitive processes [1–4]. *Paramecium* in particular is a large unicellular eukaryote organism (100 – 300 *μ*m long) whose entire surface is covered with thousands of cilia that beat synchronously [5]. When it hits an obstacle, the mechanosensitive channels at its front open and trigger an avoiding reaction: *Paramecium* swims backwards for a short time, then reorients and swims forward in a new direction [6]. On the contrary, when *Paramecium* is poked on its posterior end, it displays an escape reaction during which it accelerates forward, a response that is elicited by a different type of mechanosensitive channel [3]. The behavior of *Paramecium* in complex environments is therefore dependent on its mechanosensitivity.

At the cellular level, it is known that a mechanical stimulation opens mechanosensitive channels located in the plasma membrane, which are specifically permeable to calcium [7–9]. The resulting inward calcium current depolarizes the membrane, which then opens voltagegated calcium channels in the cilia, triggering an action potential and the reversal of the cilia [10, 11]. Mechanotransduction has been previously studied using electrophysiology experiments [2, 3]. However, in these experiments, the mechanical stimulation was systematically applied with an external probe and not elicited upon swimming. Conversely, in behavioral studies, the swimming behavior of *Paramecium* in the presence of obstacles has been reported qualitatively but not systematically quantified [1], except for the special cases of microfluidic channels [12, 13], sliding along surfaces [14, 15] or during direct interactions between paramecia [16].

In this paper, we study the swimming behavior of *Paramecium* in controlled crowded environments and focus on its local interactions with obstacles. In recent years, similar approaches have been developed for a different microorganism, the microalga *Chlamydomonas reinhardtii* (*CR*), which swims through the synchronized beating of two front flagella. On long time scales, the presence of dense arrays has been shown to decrease the effective diffusivity of *CR* [17] or deflect their trajectories [18]. On short time scales, the local interactions of *CR* with either flat or curved surfaces have been studied and characterized through both contact and hydrodynamic modeling [19–21]. However, none of these studies has explicitly taken into account the mechanosensitive properties of the micro-swimmer in the modeling of their interactions with surfaces.

In our study, we distinguish between the hydrodynamic interactions and the contact regime that can lead to a mechanosensitive response. For interactions that do not lead to an avoiding reaction, we recover a behavior that has been reported previously for *CR* [20]: when *Paramecium* contacts a pillar, it is scattered with a fixed angle, while when it interacts without contact, its trajectory is linearly deflected. When an avoiding reaction is elicited upon contact, we show that the mechanosensitive behavior can be separated into two distinct responses: an avoiding reaction that is triggered immediately upon contact with the obstacle, and an avoiding reaction which is delayed. We then show that both responses can be accounted for by a simple model where a mechanotransduction current is integrated during contact until an excitability threshold is reached. The model accounts for all our observations, but in contrast to previous studies on immobilized cells stimulated by moving rods [3, 22], it predicts that with an ecological stimulus, the mechanotransduction current is small and has a persistent component.

## II. RESULTS

### A. Interactions with a pillar

Typical experiments consist in tracking *Paramecium tetraurelia* swimming in elastomer-based pools whose bottom is either smooth or decorated with cylindrical obstacles of radius *r_p_* ≃ 150 *μ*m, spatially distributed on random and square lattices with surface fractions Φ ranging from 0.011 to 0.28 (see Table 1, Materials and Methods and electronic supplementary material). As shown in Fig. 1*A*, the height of the pillars matches the depth of the pool *h* = 340 *μ*m and paramecia are thus constrained to the volume between the pillars and cannot swim above them. In addition, since the depth of the pool is about 3 times the length of *Paramecium*, the cells can swim helicoidally but their trajectories are mostly constrained in two dimensions. Some reorientation events associated with avoiding reactions can actually occur perpendicularly to the observation plane (in about 21% of the cases for *ϕ* = 0, see electronic supplementary material for details). Figure 1*B* shows two examples of such microengineered pools (a square and a random lattice) imaged with a dark field illumination, so that both the edges of the pillars and paramecia appear as bright objects on a darker background (see white arrows on both images to locate the paramecia).

**FIG. 1.**
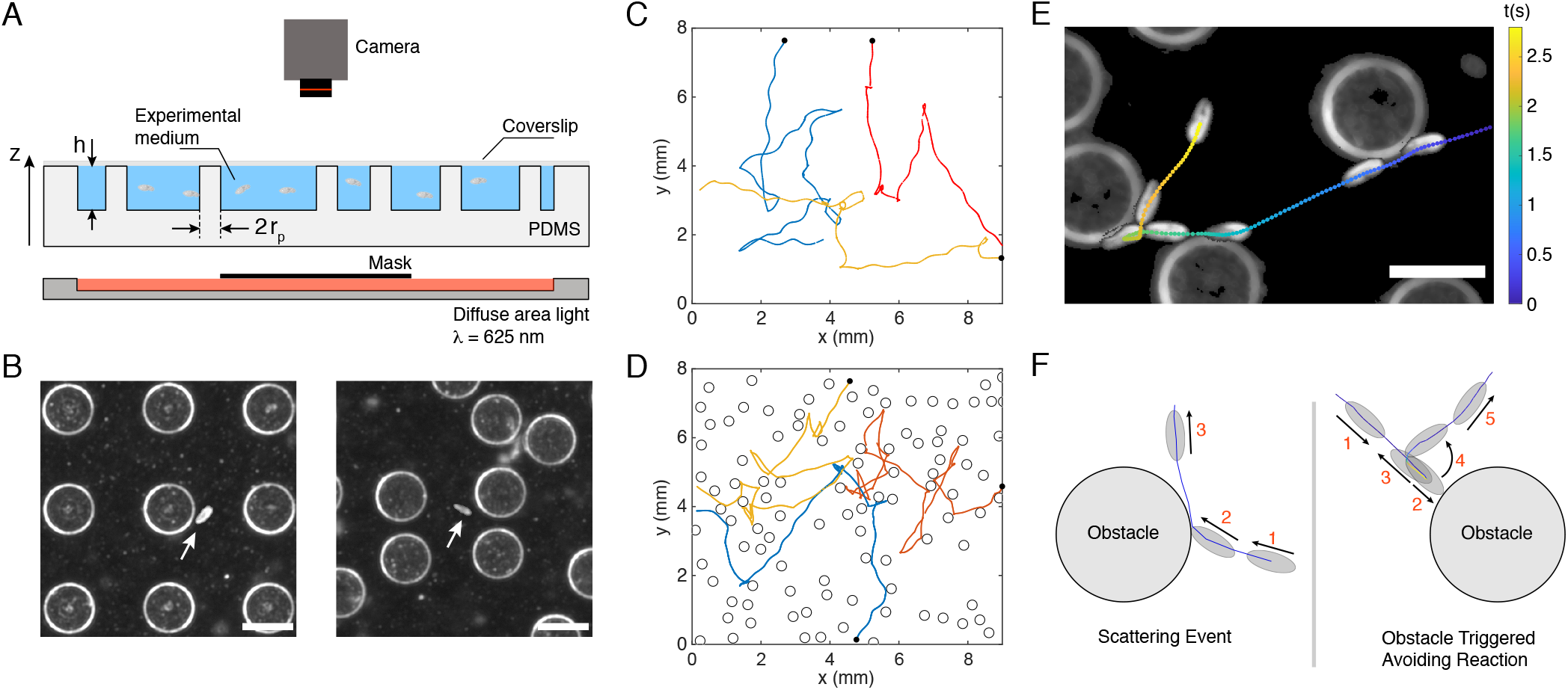
Tracking *Paramecium* in micro-engineered environments. (A) Sketch of the experimental setup (*side view*). Paramecia are deposited in a PDMS elastomer pool filled with the experimental medium (see Materials and Methods) and decorated with cylindrical pillars (height *h* = 340 *μ*m, diameter 2*r_p_* ≃ 300 *μ*m). The pool is covered with a glass coverslip to prevent evaporation. Paramecia are illuminated in dark field with a red LED panel (λ = 625 nm) and imaged from the top with a fast and sensitive camera. (*B*) Typical dark-field images of paramecia (white arrows) swimming in a square lattice (*left panel*, surface packing fraction Φ = 0.2) and in a random lattice (*right panel*, Φ = 0.083). The scale bar is 300 *μ*m long. (*C, D*) 2D trajectories of 3 paramecia in a pool, (C) without obstacles and (D) with obstacles (Φ = 0.28, random). For each trajectory, the black disk marks its starting point. (*E*) Composite image showing a close up view of the trajectory of a paramecium swimming in a random lattice of obstacles (Φ = 0.14). The color of the trajectory codes for the elapsed time and a few successive positions of the same paramecium at different times are overlayed. The scale bar is 300 *μ*m long. (*F*) Sketch of the two types of interactions with a pillar: (*left*) Scattering Event (*SE*), (*right*) Obstacle Triggered Avoiding Reaction event (*OTAR*). On both sketches, the black arrows indicate the direction of motion and the numbers the successive steps of the events.

**TABLE I.** Environments used in this work. The first column provides the lattice type. The other ones give the pillar surface fraction Φ and number *M* of experiments performed (brackets). In addition, 10 experiments were done without pillars (Φ = 0) to observe the free swimming of the cells.

|  |  |  |  |  |
| --- | --- | --- | --- | --- |
| square | 0.083 (8) | 0.14 (13) | 0.20 (9) | 0.28 (9) |
| random | 0.011 (13) | 0.083 (6) | 0.14 (10) | 0.28 (7) |

Without obstacles, the normal motion of *Paramecium* alternates between helicoidal swimming runs and spontaneous reorientations as shown in Fig. 1*C* with the example of three typical trajectories. These events have been coined “avoiding reactions” (*AR*) as they usually occur in response to different types of stimuli, such as a local change of temperature or the presence of attractants and/or repellents in the environment [1, 6]. During an *AR*, *Paramecium* comes to a stop, swims backward for a short period of time and then performs a 3D rotation of its body around its posterior end, before resuming its forward swimming motion in a new direction (see electronic supplementary material for the detailed dynamics of the *AR*, Section III, Fig. S3 and Movie S1). It results at long times in a motion (Fig. 1*C*) that is thus reminiscent of the classical *run and tumble* swimming dynamics of *E. Coli* bacteria [23].

As first reported by Jennings more than a century ago [1], an *AR* can also be triggered mechanically when *Paramecium* comes into direct contact with an obstacle, as depicted on the trajectories of Fig. 1*D*, obtained for paramecia swimming in a random network. In this case, the *AR* likely results directly from the opening of mechanosensitive channels located in the plasma membrane of *Paramecium*. However, contacts with an obstacle clearly do not only induce such triggered *AR*, but also lead to reorientation events during which *Paramecium* is “passively” scattered by the obstacle. Figure 1*E* shows a typical trajectory for which *Paramecium* first slides against two pillars at early and intermediate times (in blue), exhibiting passive-like scattering events (see electronic supplemental material, Section V, Fig. S4 and Movie S2), and then performs an *AR* upon hitting a third pillar (see the green to yellow colored points of the trajectory in Fig. 1*E*). For the rest of the manuscript, we will refer to the first two events of Fig. 1*E* as Scattering Events (or *SE*) and to the last one as an Obstacle Triggered Avoiding Reaction (or *OTAR*). Both types of events are sketched in Fig. 1*F*.

### B. Passive interactions

How does *Paramecium* passively interact with an obstacle? To quantify this, we have followed the same approach as Kantsler *et al.* [19] and Contino *et al*. [20] by studying how the cells interact with a pillar depending on their incident angle, in the absence of any *AR*. For this purpose, we define an interaction corona as the circular region of radius *r_int_* = 256 *μ*m and centered on the pillar, thus yielding a ring of width about one cell length around the pillar. We denote *θ_i_* (*resp. θ_o_*) the angle between the local radial direction and the orientation of the cell when it enters (*resp*. leaves) the interaction corona (Fig. 2*A*, inset).

**FIG. 2.**
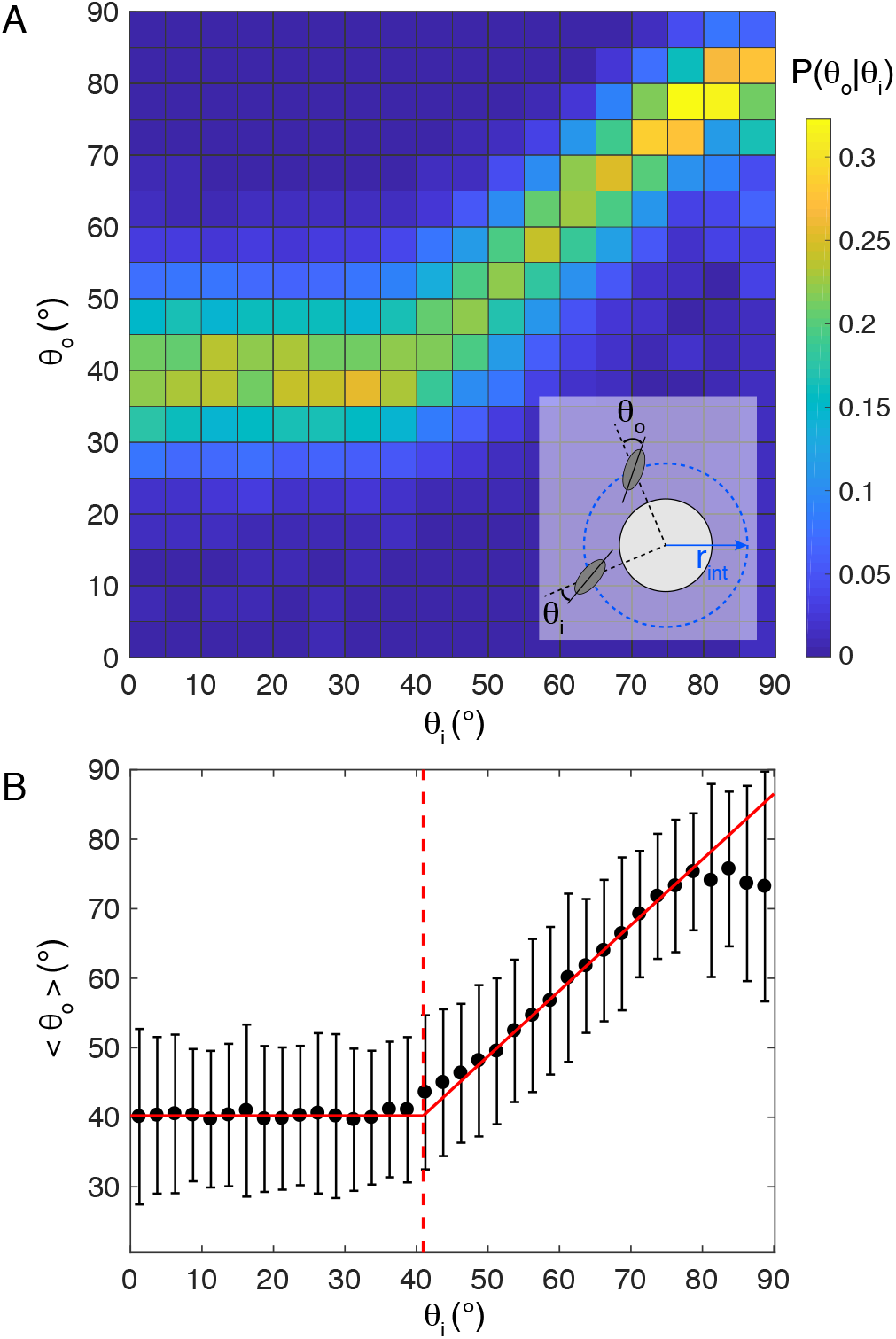
Characterizing passive interactions. (*A*) Conditional probability *P*(*θ_o_*|*θ_i_*) for *θ_i_* and *θ_o_* varying between 0° and 90°. Each square bin has a side length of 5°. This plot was obtained from *N* = 28031 events compiled from *M* = 75 independent experiments ranging from *ϕ* = 0.01 to *ϕ* = 0.28 in a square or random lattice. *Inset*: sketch to define the geometrical parameters. The blue dashed circle corresponds to the interaction corona while the solid black circle corresponds to the pillar. A paramecium intercepts the interaction corona with an angle *θ_i_* and leaves it with the angle *θ_o_*. (*B*) Mean output angle 〈θ_o_〉 versus *θ_i_*, where *θ_i_* is taken as the center value of each bin, of width 2.5°. Solid red lines are linear fits of the contact and hydrodynamic regimes, that intercept at 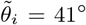, marked by the vertical red dashed line. In the hydrodynamic regime, the linear fit yields, for *θ_i_* ∈ [50°, 80°], a slope *m* = 0.94 ±0.031 and intercept *q* =1.6 ± 2.0° with *R*^2^ = 0.99.

We have measured *θ_i_* and *θ_o_* over *N* = 28031 passive interaction events, *i.e*. events during which the cell has traveled through the interaction corona without any *AR*, recorded from *M* = 75 independent experiments (Fig. 2*A*). Below a threshold incidence angle 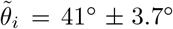, cells always leave the pillars with the same mean angle 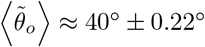. In this case, there is no memory of the in-going angle *θ_i_*. Above 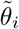, the cell travels through the interaction corona without actually contacting the pillar. In that regime, its trajectory is simply deflected and the mean angle 〈*θ_o_*〉 depends linearly on *θ_i_* with a slope *m* = 0.94 ± 0.031 (Fig.2*B*). The measured value of 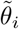 matches a simple geometric prediction, based on a tangent contact of the cell with the pillar, 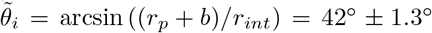 with the semi-width of the cell *b* = 29.5 ± 4.4 *μ*m.

The measurements for the contact regime are coherent with our observations that upon contact, the cell first reorients and aligns along the pillar, then slides against it before leaving tangentially to the pillar, independently of the in-going angle. Finally, the conditional probability map *P*(*θ_o_*|*_i_*) found for *Paramecium* is very similar to the one measured by Contino *et al*. [20] with the biflagellate microalga *CR*, suggesting a universal behavior at contact among microswimmers, independently of their swimming mode. In the following, we will focus exclusively on the interactions during which the cell collides mechanically with the obstacle and we will thus not consider the hydrodynamic regime.

### C. Obstacle triggered avoiding reactions

Although it is well known that mechanical stimulation of the front part of *Paramecium* can trigger an *AR* [1, 24], it is unclear how *ARs* are elicited upon contact on larger obstacles such as our pillars, and at the swimming speed of *Paramecium* itself. We first ensured that *ARs* in the vicinity of obstacles were indeed triggered by the mechanical contact with the obstacle, *i.e*. that they are *OTARs* and not spontaneous *ARs* happening close to it by chance. To do so, we looked at the radial dependence of the *AR* frequency *f_AR_*(*r*) defined as

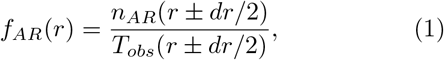

where *n_AR_* is the total number of *AR* in the annulus of width *dr* at position *r*, and *T_obs_* is the total observation time of cells in this annulus (see inset in Fig. 3).

**FIG. 3.**
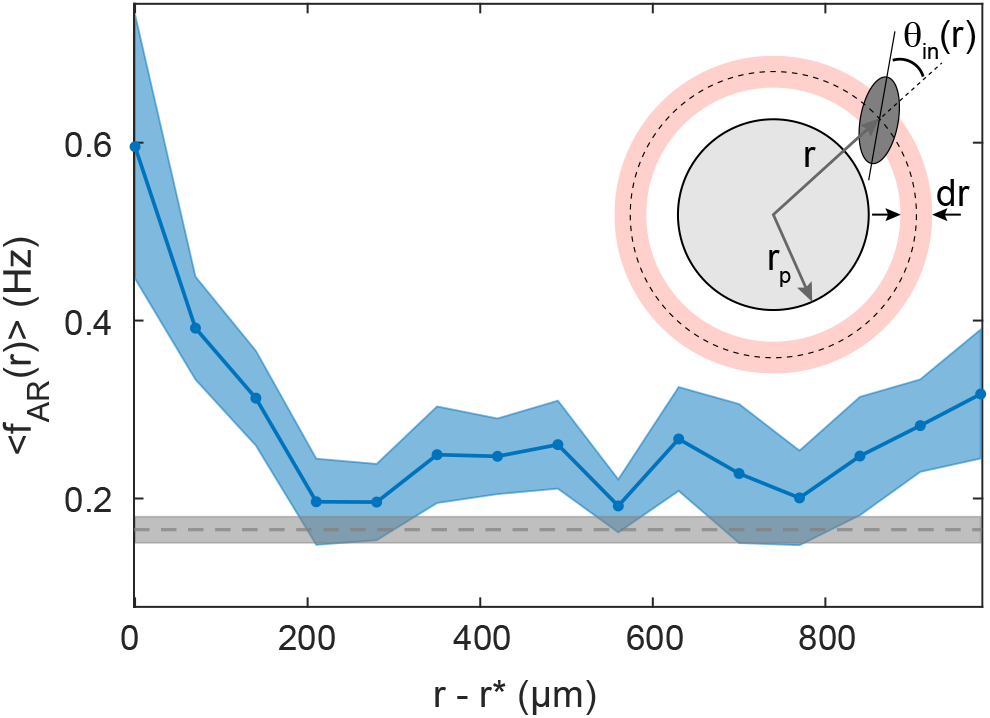
Detecting *AR*s in the vicinity of a pillar. Average *AR* frequency 〈*f_AR_*(*r*)〉 at a distance *r* – *r*\* from the surface of the pillar. The blue curve is obtained by averaging *f_AR_*(*r*) from *M* = 13 independent experiments and the blue shaded area corresponds to the standard error of the mean (*SEM*). The experiments were performed in a randomly distributed pillar array with Φ = 0.01. The horizontal gray dashed line corresponds to the mean avoiding reaction frequency 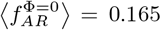 Hz in the absence of obstacles and computed over *M* = 10 independent experiments, and the gray shaded area gives the *SEM*. Inset: sketch to illustrate the geometrical parameters. For these measurements, a bin size *dr* = 70 *μ*m was chosen, that matches the average semi-length of a cell.

Figure 3 shows 〈*f_AR_*(*r*)〉 plotted as a function of the distance from the surface of the pillar *r* – *r*\*, with *r*\* = *r_p_* + *b* ~ 172 *μ*m the inaccessible volume radius, and where the brackets denote an averaging over *M* = 13 independent experiments. In the immediate vicinity of a pillar, one clearly sees that the spatial *AR* frequency increases by a factor of about 4 compared to its value far from the pillar, peaking at *r*\*. Such an increase is a clear signature of a mechanosensitive response. In addition, far from the pillar, one recovers the mean value of the spontaneous *AR* frequency measured in pillar-free environments, 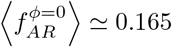 Hz.

Direct visual inspection of paramecia swimming against pillars reveals that not all*OTAR* are actually triggered instantaneously upon contact (see electronic supplemental material, Movies S3 and S4). To quantify this effect, we have measured the duration between the collision time and the start of the backward swimming that marks the beginning of an *OTAR* (see electronic supplemental material and Materials and Methods for the determination of both times). This duration, noted *τ*, is called the triggering time. Figure 4*A* shows the distribution of *τ* obtained with a total of *N* = 1790 *OTAR*s. This distribution has two peaks, a first narrow one at *τ* ≈0 s and a second broader one at *τ* ≈ 0.15 s. Thus, *Paramecium* displays two types of reactions: an immediate one, referred to as an *instantaneous OTAR* (*τ* ∈ [0, 0.04] s), and a delayed one, called *delayed OTAR* (*τ* ∈]0.04, 0.6] s).

**FIG. 4.**
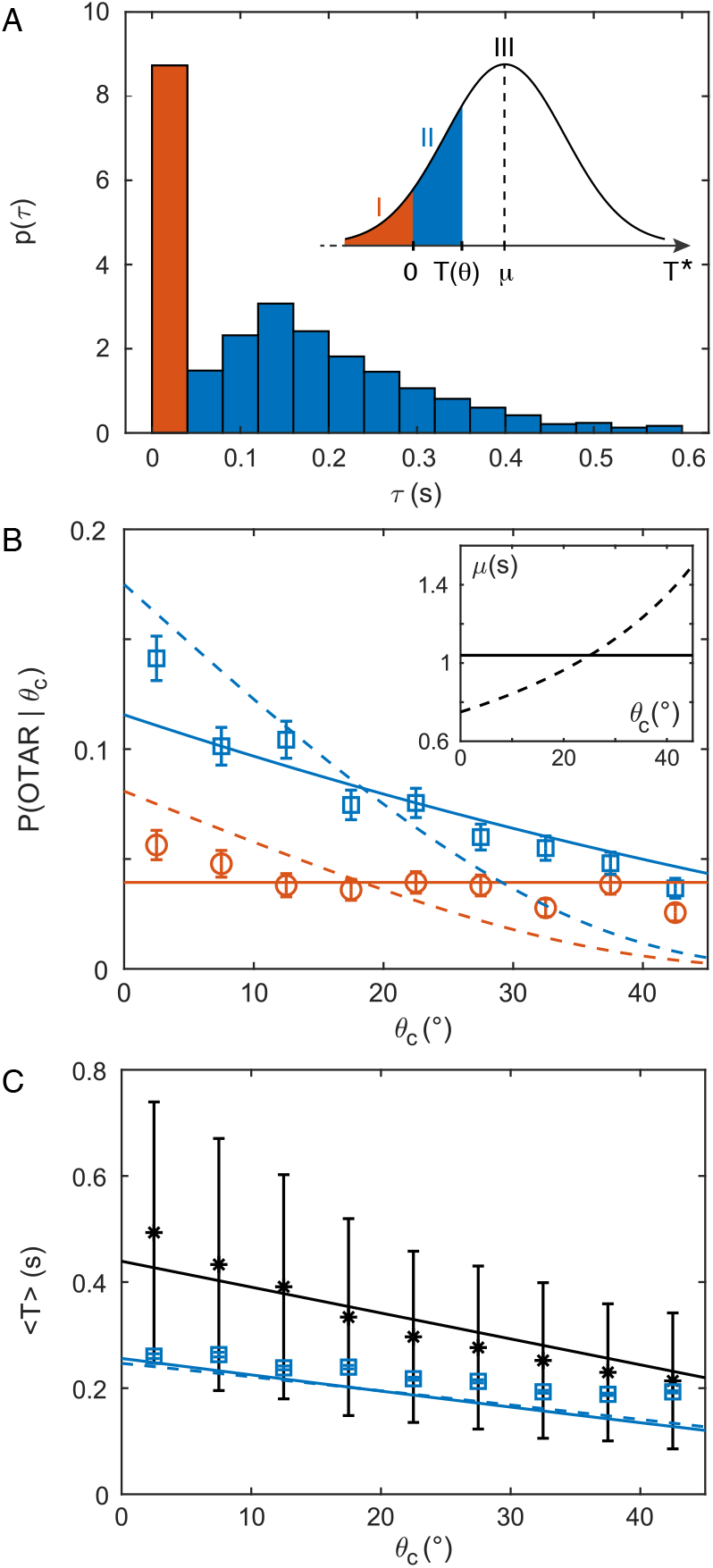
Identification and modeling of two types of *OTAR*s. (*A*) Probability density function of the measured triggering time *τ* with a bin width of 2 frames. Red and blue bars correspond respectively to instantaneous and delayed *OTAR*s. Inset: sketch of the model distribution of *T*\* with mean μ, along with the three modes. (*B*) Probability of doing an instantaneous (red circles) or a delayed (blue squares) *OTAR*given the collision angle *θ_c_*, as a function of *θ_c_* with a bin width Δ*θ_c_* = 5° (see Materials and Methods for the error bar calculation). The solid (*resp.* dashed) line is for the uniform (*resp*. non-uniform) model with constant *μ*(*resp*. *μ*(*θ_c_*) = *c*/(90 – *θ_c_*)). Inset: mean *μ* of the triggering time *T*\* as a function of *θ_c_* for the uniform response model (solid line) and non-uniform response model (dashed line). (C) Average contact duration 〈*T*(*θ_c_*)〉 for *SE*s (black stars) and for delayed *OTARs* (blue squares). The solid black line is a linear fit of the *SE* data using equation (2), yielding *T_max_* = 0.44 s (with *R*^2^ = 0.87). The solid (*resp*. dashed) blue line corresponds to the prediction of contact duration in the case of delayed *OTARs*, with constant *μ* (*resp*. depending on *θ_c_*).

Furthermore, we find that the probability of an instantaneous *OTAR* does not depend on the contact angle *θ_c_* (defined as the incident angle *θ_in_* at contact, see inset of Fig. 3 and Materials and Methods), while the probability of a delayed *OTAR* decreases with increasing *θ_c_* (Fig.4*B*). We present below an electrophysiology-based model that accounts for these two distinct responses.

### D. A simple model

How can *Paramecium* display two different types of behavior upon contact, with a different dependence on the incidence angle? Electrophysiological studies on immobilized cells show that a mechanical stimulation of the front membrane triggers an inward calcium current with short latency and duration, in the millisecond range [3, 22]. This transient current then depolarizes the membrane and quickly triggers an action potential, initiating the avoiding reaction. Given this evidence, the observation of *OTAR*s delayed by several hundreds of milliseconds is surprising. We propose here a simple model to explain both types of reactions. First, given that most contacts do not trigger an avoiding reaction, we postulate that the current triggered upon contact is smaller than in typical mechanical stimulations of immobilized cells. To trigger an action potential, the charge *Q* transmitted at contact must exceed a threshold *Q*\* ≈ *CV*\*, where *C* ≈ 300 pF is the membrane capacitance [11] and *V*\* ≈ 3 mV is the threshold potential to trigger an action potential relative to the initial potential, which quantifies cell excitability [25]. Thus, *Q*\* ≈ 1 pC. Depending on the cell excitability at the instant of contact, the transmitted charge may or may not trigger an action potential.

Second, to account for the delayed reactions, we postulate that, in addition to the instantaneously transmitted charge, there is a small transduction current *I*_0_ that persists as long as the cell is in contact with the pillar. This could be due to an incomplete inactivation of the mechanosensitive channels, as observed in patch clamp recordings of *Piezo* channels [26], or to the progressive recruitment of mechanosensitive channels during sliding, as different parts of the membrane come into contact with the pillar. Therefore, during contact, a transmitted charge *Q* + *I*_0_*T* accumulates during a time *T* until it reaches the threshold *Q*\* or the cell leaves the pillar. Thus, an avoiding reaction is triggered if the cell remains in contact for a minimum duration *T*\* = (*Q*\* –*Q*)/*I*_0_. We assume that *T*\* is a random variable normally distributed with probability density *p*, mean *μ* and standard deviation *σ*. This would occur for example if the membrane potential (which is noisy [27]) was normally distributed. Thus, we can see that three cases can occur: *(i)* for *T*\* ≤ 0, a contact instantaneously triggers an avoiding reaction: this is an instantaneous *OTAR; (ii)* the cell stays in contact for a time *T*\*, then does an avoiding reaction: this is a delayed *OTAR*; *(iii)* the cell leaves the obstacle before time *T*\*: this is a scattering event *SE*.

To quantify the probability of each case, we first measured the duration *T*(*θ_c_*) of a *SE* as a function of the contact angle *θ_c_* (Fig. 4*C*, black stars). If we assume, as our observations suggest, that the cell slides on the pillar until it leaves it tangentially, then this duration should depend approximately linearly on *θ_c_* as follows:

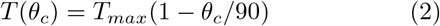

Fitting this expression to our data (Fig. 4*C*, black solid line) yields *T_max_* = 0.44 ± 0.035 s (95% confidence bounds, *R*^2^ = 0.87). Note that the same trend for *T*(*θ_c_*) was measured for *Pawn* cells, a mutant of *Paramecium tetraurelia*, which cannot perform *AR*s due to a lack of voltage-gated calcium channels in the cilia [28] (see electronic supplementary material, Section VIII, Fig. S7). This confirms the passive nature of *SE*s.

We can now express the probability of each of the three cases (Fig. 4*A*, inset).

**Case 1:** An instantaneous *OTAR* occurs with probability

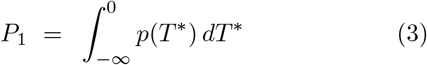

**Case 2:** A delayed *OTAR* occurs with probability

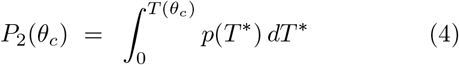

**Case 3:** A *SE* occurs with probability

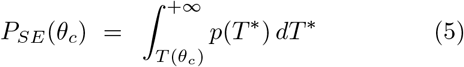

The model has two free parameters, *μ* and *σ*, which we constrain with two experimental data points, *P*_1_(*θ_c_* = 22.5°) and *P*_2_(*θ_c_* = 22.5°). We find *μ* ≈ 1.04s and *σ* ≈ 0.59 s, which means that it takes on average about 1 s for a contact to trigger an avoiding reaction.

The model predicts that the probability of an instantaneous *OTAR* does not depend on *θ*c**, in agreement with our observations (Fig. 4*B*, red solid line), because the transduction current is assumed to be independent of the contact position. Less intuitively, the model also predicts that the probability of a delayed *OTAR* decreases with increasing *θ_c_*, because contacts with smaller *θ_c_* are longer. This angular dependence is in quantitative agreement with our observations (Fig. 4*B*, blue solid line).

Electrophysiological studies show that the transduction current vanishes in the middle of the cell (*θ_c_* = 90°) (and reverts on the posterior side) [29]. What would happen if it decreased gradually over the anterior part with increasing *θ_c_*? We calculate the probabilities of the three cases with an angular dependence of the mean of *T*\*, as follows:

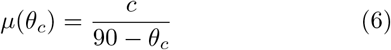

where the constant *c* is chosen such that the angular average 〈*μ*(*θ_c_*)〉 equals the mean of *T*\* in the uniform model (Fig. 4*B*, inset). Here, the triggering time *T*^*^ is minimum at θ_c_ = 0°and infinite at *θ_c_* = 90°. With this model, we find that the probabilities of both instantaneous and delayed *OTAR*s vary substantially with the contact angle *θ_c_*, unlike in the observations (Fig. 4*B*, red and blue dashed lines).

Finally, the model allows predicting the mean contact duration of delayed *OTAR*s as a function of *θ_c_*, which is

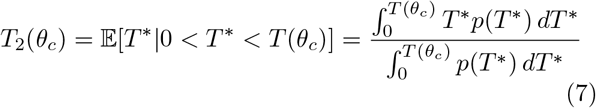

The prediction is in quantitative agreement with the observations (Fig. 4*C*, blue solid and dashed lines). These values are in the range of the measured secondary peak of the distribution of the triggering times *τ* (Fig. 4*A*). Even though the mean charging time is about 1 s, the triggering time is in fact limited by the maximum sliding duration against the pillars (about 0.3s, Fig. 4*C*, blue squares).

## III. DISCUSSION

Motile microorganisms naturally evolve in complex environments that they have to sense and react to [30]. It is therefore critical to understand how their foraging efficiency depends on their local interactions with obstacles found in their natural habitat. In order to answer these questions, a large array of literature has been dedicated to the study of motility in engineered complex environments in the past years [17, 20, 21, 31].

In this context, *Paramecium* is interesting as it can be studied both through behavioral and electrophysiology approaches thanks to its large size. As a matter of fact, its swimming behavior as well as its mechanosensitive response, have been extensively described in past literature, reviewed in [6]. The main message from these studies is that *Paramecium* performs an avoiding reaction upon collision with an obstacle, through a mechanosensitive response, and this response is modulated by the location of the stimulation on the surface of the swimmer. However, our work reveals a different picture for the behavior of this microorganism in our experimental conditions.

First of all, collision with an obstacle does not necessarily elicit an avoiding reaction. In fact, it is passively scattered on the surface of the pillars 90% of the time, which means that its trajectory is deflected without exhibiting any backwards swimming. In this case, the scattering event imposes the angle *θ_o_* at which the swimmer escapes the immediate neighborhood of the pillar. Conversely, when *Paramecium* interacts hydrodynamicaly with the pillar, this angle *θ_o_* depends linearly on the incident angle *θ_i_*. In the absence of an avoiding reaction, our findings directly echo previous observations on a different microorganism [20], the microalgae *Chlamydomonas reinhardtii*.

The fact that only 10% of the contact events lead to an avoiding reaction is surprising. Actually, a similar small rate of avoiding reactions (of less than 5%) has been reported by Ishikawa *et al*. [16], although in a different context that involves collisions between two freely swimming paramecia. Apart from this particular case, such an observation has not been quantified in previous behavioral experiments of freely swimming paramecia, which were done more than a century ago [1]. Electrophysiological studies on immobilized cells on the other hand, did not suggest however such a low rate of mechanical reactions [3, 8, 9, 22]. This may be due simply to the fact that mechanical stimulation with probes was adjusted precisely so as to induce a measurable mechanosensitive response (see in particular Fig. 3 in [3]). To our knowledge, all electrophysiological studies on the mechanotransduction of *Paramecium* used micrometer-sized glass probes, and it could well be that the applied force were much stronger than in our experiments, especially as our obstacles are much wider than typical probes.

From a behavioral perspective, it is intriguing that avoiding reactions are relatively rare when *Paramecium* hits large obstacles such as our pillars. One possible explanation is that it might not be critical for the organism to react to a large obstacle when it can be passed by sliding. This would explain the occurrence of delayed reactions: an avoiding reaction is triggered when the organism is blocked by the obstacle for a certain amount of time. Another possibility could be that mechanosensitivity serves not simply navigation in crowded environments, but also and perhaps primarily the avoidance of sharp objects that may harm the membrane, as is suggested by evolutionary accounts [32].

Another surprising finding is that many *OTAR*s are delayed by about 150 ms, when electrophysiological studies report nearly instantaneous responses. According to voltage clamp measurements, mechanotransduction currents activate with a very short latency and a rise time smaller than 20 ms, in *Paramecium* and other ciliates [8]. In addition, the current appears to be transient. There are two possibilities to make our findings consistent with these previous studies. One is that the current inactivates only partially. Indeed, this is what has been observed in Piezo channels with a constant applied pressure [26]. Previous studies in *Paramecium* quantified the peak current but not its stationary value. Another possibility is that, as the organism slides along the obstacle, additional mechanosensitive channels are recruited.

Finally, another noticeable discrepancy with previous work is the distribution of mechanosensitivity along the body of *Paramecium*. While our experimental data is in agreement with a homogeneous distribution of mechanosensitivity along the front part of *Paramecium*, previous electrophysiological studies showed graded responses along the body axis [22]. Stimulating the anterior part triggers an inward current, carried mostly by calcium, while stimulating the posterior part triggers an outward current, carried by potassium. In the middle, the transduction current vanishes. When potassium channels are blocked pharmacologically, mechanical stimulation triggers a uniform depolarizing response across the body [3]. Thus, the spatial gradient of mechanosensitivity appears to be due to the cancellation of a spatially uniform depolarizing mechanoreceptor current by a spatially graded hyperpolarizing current. It is not entirely clear from previous measurements whether this cancellation is linear between the anterior and posterior end, and this is complicated by experimental difficulties. Indeed, immobilized cells were mechanically stimulated with a glass probe of fixed orientation, and with a fixed movement, while position along the membrane was varied. Thus, the orientation and direction relative to the membrane surface varied, and the amplitude of the membrane deflection was not fixed either. In contrast, in our experiments, the obstacles are very controlled in their shape and mechanical properties. Therefore the contact only depends on the swimming behavior of *Paramecium* itself. Moreover, it is unclear how the presence of electrophysiology micropipettes inside the organism and its immobilization might affect its mechanical properties. For instance, the presence of a pipette might impose a prestress on the cell surface. In fact, in artificial mechanoreceptors, wetting of the membrane on a pipette is sufficient to trigger the opening of mechanosensitive channels [33].

In the future, observing electrophysiological responses of swimming organisms to obstacles should therefore be key to unify findings obtained from electrophysiology and behavioral experiments.

## IV. MATERIALS AND METHODS

### A. Cell culture and preparation

Experiments were performed with the wild-type stock 51 of *Paramecium tetraurelia*, obtained from Eric Meyer, Institut de Biologie, Ecole Normale Supérieure, Paris, France. The cells were cultured in a fully opaque incubator set at 27°C in a buffered infusion of wheat grass powder (Wheat Grass Powder, Pines) complemented with 0.8 *μ*g/mL β-sitosterol and inoculated with non-pathogenic *Klebsiella pneumoniae* as a food source. After 48 h of growth and at least 20 min before an experiment, about 0.4 mL of cell suspension were pipetted from the surface of the culture medium into 4 mL of an inorganic medium (1mM CaCl_2_, 4mM KCl and 1mM Tris HCl, pH 7.2). As a control, experiments with the *Pawn* mutant strain, which cannot perform *AR*s due to a lack of voltage-gated calcium channels in the cilia [28], were also performed. The same culture and preparation protocols were carried out.

### B. Fabrication of the pools

Pools were manufactured using a combination of micro-milling and elastomer molding techniques fully described in the electronic supplementary material, Section I and in Fig. S1. They are made of a polydimethylsiloxane elastomer (PDMS; Sylgard 184, Dow Corning; cross-linking ratio 10:1, Young’s modulus *E* ≃ 2.7 ± 0.8 MPa) and consist of a square wall of height *h* = 340 *μ*m and edge length 30 mm that delimits an accessible volume. Its surface is either bare or decorated with cylindrical pillars of radius *r_p_* = 142.5 *μ*m, distributed according to a square lattice or randomly. For the square lattice, the surface fraction 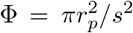 is varied by changing the mesh size *s*. For the random lattices, pillars are distributed randomly with a minimal spacing of ≃ 60 *μ*m to avoid trapping of the cells. Prior to an experiment, the elastomer pool was exposed to an oxygen plasma for about 1 min to render the PDMS surface hydrophilic. After injecting the cells, the pool was closed with a glass coverslip in contact with the top of the wall and the pillars.

### C. Experimental setup and imaging

Cells were imaged from the top with a variable zoom lens (MVL12X12Z, Thorlabs) set on 1.5× combined with a 1.33x extension tube (Thorlabs, MVL133A) yielding a pixel size of 3.81 ±0.01 *μ*m. Images were captured with a high resolution and sensitive CMOS camera (Black-fly S BFS-U3-51S5M-C, Flir, USA, 2448 × 2048 pixels^2^, 10 bits) operating at 50 fps, using its dedicated acquisition software *Spin View*. To enhance contrast, the pool was illuminated in a dark field configuration using a square LED panel (EFFI-SBL, Effilux, France) as a light source, producing a red light (λ = 625 nm) to minimize phototaxis [34].

### D. Image processing and analysis

Raw images were directly recorded to a hard drive and compressed by first removing the background then applying a threshold [11]. Automatic tracking was performed on compressed images using *FastTrack* [35]. To maximize tracking quality, tracked movies were then visually inspected and the few remaining errors were manually corrected using the embedded post-processing tools of the *FastTrack* program. At each time t, the cell contour was fitted by an ellipse whose orientation *θ*(*t*), defined as the angle between the major axis of the ellipse and the horizontal axis *x* of the images (see Fig. 1*C*, *D* and electronic supplementary material, Section II, Fig. S2), was computed from the asymmetry of the pixel histogram along the major axis. The semi-major and semi-minor axes were estimated at *a* = 68.5±7.7 *μ*m and *b* = 29.5±4.4 *μ*m, respectively (average±std, *N* = 5080 trajectories).

### E. Post-processing

A *Matlab* (Mathworks Inc. USA) script was used to remove trajectories shorter than 1 s and time intervals during which the object was immobile. Another *Matlab* script was used to remove trajectories that presented circling motions, based on a measure of the sign of the instantaneous angular velocity 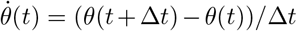 with Δ*t* = 20 ms, the time interval between frames.

### F. Detection of avoiding reactions

In our analysis, an avoiding reaction (*AR*) is described as a two-step process: a backward swimming (*BS*) followed by a reorientation. The *BS* was detected when the instantaneous motion vector **m**(*t*) and the orientation vector **o**(*t*) (posterior to anterior) pointed to opposite directions, *i.e*. **m**(*t*) o **o**(*t*) < 0. *BS* events consisting of a single frame were discarded. The reorientation event was identified as the time interval during which the instantaneous angular speed was large enough, *i.e*. 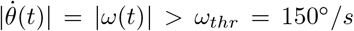 (see electronic supplementary material, Section III and Fig. S3).

### G. Detection of mechanical contacts

We defined a contact corona of radius *r_c_* = *r_p_* + *δr* where *δr* = 19 *μ*m was chosen using physical arguments and visual inspections (see electronic supplementary material for details, Section VI and Fig. S5*A*). This sets a contact interaction interval 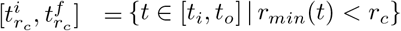, where *r_min_*(*t*) is the distance between the center of the pillar and the ellipse that fits the cell at time *t*, and where *t_i_* (*resp. t_o_*) is the time at which the cell center enters (*resp*. leaves) the interaction corona. We considered that the cell collided with the obstacle if and only if its incident angle *θ_in_* when entering the contact corona was smaller than a threshold, namely 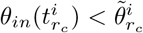, where the contact angle threshold is defined in the same way as for 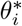 but at 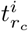. (see electronic supplementary material for details, Section VI and Fig. S5*B,C*). In this case, the collision time 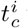, was defined as the first velocity minimum within the contact interaction interval 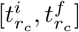 defined above. The interaction was categorized as a *SE* if no *AR* was triggered within the contact corona. In this case, the end of a contact was defined as the first time at which the cell is tangent to the obstacle, *i.e*. 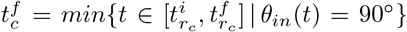. An *AR* triggered within the contact corona may be an *OTAR*, or a spontaneous *AR* occurring shortly after a *SE*. We used the latency of the *AR* to determine more finely whether this was an *OTAR* or a *SE* (see electronic supplementary material, Section VII and Fig. S6). If it was categorized as an *OTAR*, the end of the contact was the starting time of the avoiding reaction. Applying this procedure over all environments allowed us to identify a total of *N_SE_* = 15601 *SE*s, *N_inst_* = 670 instantaneous *OTARs* (*τ* ≤ 40 ms) and *N_delayed_* = 1120 delayed *OTARs* (*τ* > 40 ms).

### H. Error estimation of the conditional *OTAR* probability

We denote by *n_inst_* the number of instantaneous *OTARs* with collision angle *θ_c_* ±*dθ_c_* and *n_c_* the total number of collisions with *θ_c_* ± *dθ_c_*. Then, the number of instantaneous *OTARs* can be modeled as *n_c_* draws of a binomial distribution of parameter *p* = *n_inst_*/*n_c_* and variance mp(1 – *p*). Thus, the conditional probability *p* = *P*(*OTAR*|*θ_c_*) represented in Fig. 4*B* can be estimated as *n_inst_*/*n_c_* and its error is given by 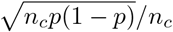. The same estimation was applied for delayed *OTAR*s.

### I. Numerical computations

Data analysis and numerical computations were done with *Matlab*. The contact duration *T*(*θ_c_*) was fitted using the Curve Fitting Toolbox with the Nonlinear Least Square method. The set of non-linear equations for *P*_1_ and *P*_2_(*θ_c_*), *i.e*. equations (3) and (4), was solved using the function *fsolve.m*.

## ACKNOWLEDGMENTS

The authors wish to acknowledge financial support from CNRS (MITI, Défi Mécanobiologie, Project PERCÉE), Sorbonne Université (Emergence-Project NEUROSWIM), Agence Nationale de la Recherche (ANR-20-CE30-0025-01 and ANR-21-CE16-0013-02), Programme Investissements d’Avenir IHU FOReSIGHT (Grant ANR-18-IAHU-01) and Fondation Pour l’Audition (Grant FPA RD-2017-2). The authors thank Eric Meyer for providing us with the wild-type of *Paramecium tetraurelia* and for insightful discussions. The authors also thank Mireille Betermier and Anne-Marie Tassin for providing us with the *Pawn* mutant of *Paramecium tetraurelia*. Finally, the authors thank Georges Debrégeas and Anke Lindner for fruitful discussions.

