## Supplementary Material for "Interaction of the mechanosensitive microswimmer *Paramecium* with obstacles"

(Dated: December 16, 2022)

This supplementary material provides additional details of the materials and methods used in this work as well as a description of the movies of the different types of interactions *Paramecium* has with a pillar.

### I. FABRICATION OF THE POOLS

Quasi-2D environments consist of square elastomer pools, the bottom of which is either bare or decorated with cylindrical pillars. These pools are obtained using a combination of micro-milling and elastomer molding techniques (Fig. S1). A Plexiglas mold that has the shape of a square mesa-like structure (side length 30 mm) is first milled with a square end mill of diameter 1 mm, using a CNC micro-milling machine (Minitech, Machinary Corp., USA). For pillar-decorated pools, the top surface of the mesa is then drilled with cylindrical wells of radius 150 $\mu$m using a micro-drilling bit. Since micro-drill bits have a conical ending, wells are then further milled with a square end mill to have a full depth $h = 340\,\mu$m. Wells are distributed with a spatial resolution of 1 $\mu$m either on a square lattice of mesh size (shortest center to center distance) $s$ or randomly, with varying surface fractions of pillars $\Phi$. For the square lattices, $\Phi$ is defined as $\Phi = \pi {r_p}^2/s^2$. For the random lattices, wells are distributed with a non-overlapping constraint and a minimal spacing between wells of about 60 $\mu$m to avoid trapping of the cells and with the same surface fraction values $\Phi$ as for the square lattices.

Pools are made of a commercial transparent elastomer, Poly-DiMethyl Siloxane (PDMS, Sylgard 184 Dow Corning, USA). They are obtained by pouring onto the Plexiglas mold a 10:1 mass ratio PDMS/crosslinker liquid mixture, that is first centrifuged at 3000 g for 5 min and put into a vacuum chamber for at least 1 hour to remove any air bubbles. The mold is then closed with a lid to ensure flatness of the bottom of the pool, and placed in an oven at 65°C for at least 12 h. The polymerized pool is then removed gently to avoid breaking the pillars formed in the wells. Before an experiment, the pool is exposed to an oxygen plasma for about 1 min to render the PDMS surface hydrophilic. This step prevents any air bubble trapping at the base of the pillars once the paramecia seeded liquid is poured into the pool. The average pillar radius was measured using an optical microscope and found to be $r_p = 142.5\,\mu$m.

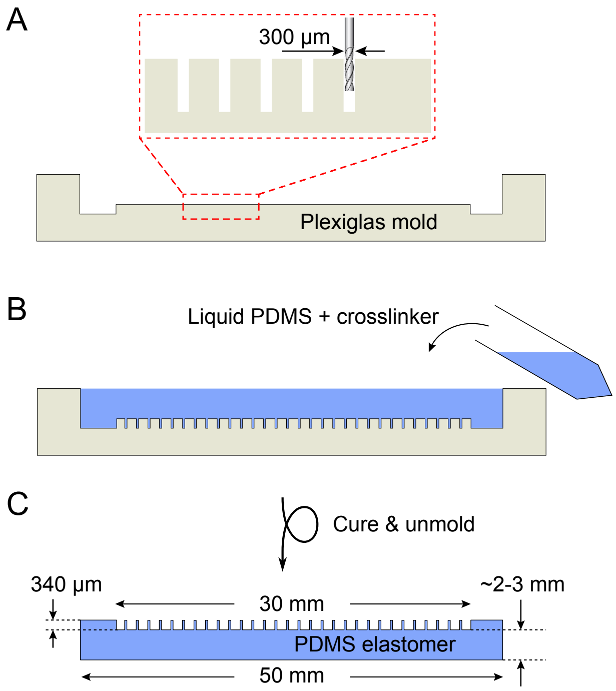


FIG. S1. Fabrication of the pools (side view) *(A)* A Plexiglas mold is micro-milled with cylindrical wells. *(B)* A liquid PDMS-crosslinker is then poured into the mold. *(C)* After curing and unmolding, one obtains a pool with pillars arranged with the desired pattern.

### II. DEFINITION OF THE GEOMETRICAL PARAMETERS

As explained in the Materials and Methods section of the main text, contours of the cells were fitted with an ellipse of semi-major and semi-minor axes $a$ and $b$ respectively, and orientation $\theta(t)$ with respect to the horizontal axis $x$ of the recorded images. Figure S2 sketches the different geometrical parameters that were used to describe the orientation of the cell in free swimming and when it interacts with a pillar of radius $r_p$.

*
†

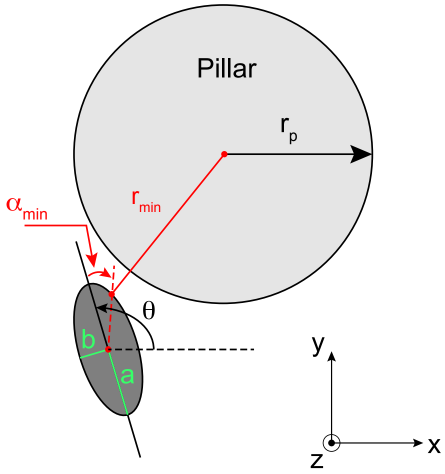


FIG. S2. Geometrical parameters used to describe the orientation of a cell in free swimming and when it interacts with a pillar. The distance between the center of the pillar and the ellipse that fits the contour of the cell is noted $r_{min}$, and the corresponding signed angle with respect to the major axis of the ellipse is $\alpha_{min}$.

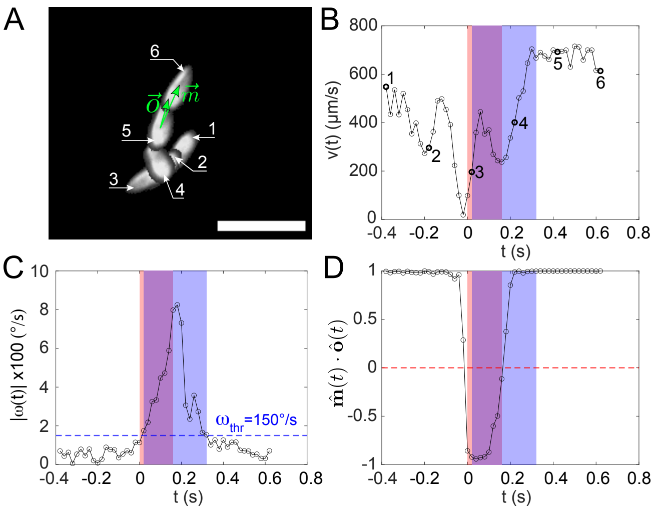


FIG. S3. Detailed dynamics of an example *AR*. *(A)* Composite dark-field image showing a close-up view of a paramecium performing a spontaneous *AR* obtained by overlapping images taken at different times separated by 200 ms. The successive positions of the cell are numbered from 1 to 6. On position 5 are overlapped the instantaneous motion vector **m** and orientation (posterior to anterior) vector **o**. The scale bar is 300 $\mu$m long. *(B-D)* Time evolution of the parameters characterizing the trajectory of the cell during the *AR* of Fig. S3*A*. *(B)* Instantaneous speed of the cell $v(t)$. The origin of time $t = 0$ is taken as the beginning of the *BS* phase. The colored regions correspond respectively to the *BS* phase only (pink), to the *BS* and reorientation phase occurring simultaneously (purple) and to the reorientation phase only (blue). Labels 1 to 6 correspond to the numbered positions of Fig. S3*A*. *(C)* Angular speed $|\omega(t)|$. The blue dashed horizontal line is the threshold for the detection of the reorientation phase. *(D)* Scalar product between the motion unit vector $\hat{\mathbf{m}}(t)$ and the orientation unit vector $\hat{\mathbf{o}}(t)$. The red dashed horizontal line is the threshold for the detection of a *BS* event. For all graphs, data points are plotted every 20 ms.

### III. DETAILED DYNAMICS OF AN AVOIDING REACTION

An avoiding reaction (*AR*) can be described as the succession of the following phases: interruption of the forward motion, backward swimming (*BS*) along a straight line, reorientation around the posterior end and forward motion. Figure S3*A* shows a composite image of a typical spontaneous *AR* occurring in a pillar-free environment and obtained by overlapping 6 consecutive dark-field snapshots separated by 200 ms. Note that in this example, the $4^{th}$ snapshot partially overlaps and hides the $3^{rd}$ snapshot.

When *Paramecium* is not doing an *AR*, it typically swims at speeds $v = 542 \pm 294\,\mu$m/s ($N = 3.58\,10^5$ data points in a pillar-free environment, $\Phi = 0$, with $M = 10$ independent experiments). These values are consistent with the measured speeds reported in Fig. S3*B* prior to an *AR* ($v \simeq 500\,\mu$m/s for $t \leq -0.1$ s, where $t = 0$ defines the beginning of the *BS*) and after the end of the reorientation phase ($v \simeq 700\,\mu$m/s for $t \geq 0.4$ s). We define the instantaneous angular velocity $\omega(t) = \dot{\theta}(t) = (\theta(t + \Delta t) - \theta(t))/\Delta t$, with $\theta(t)$ the orientation of the cell relative to the horizontal axis $x$ at time $t$ and $\Delta t$ the time interval between consecutive frames (in our case, the camera operates at 50 frames/s and $\Delta t = 20$ ms). In our experiments, we measured that overall $|\omega| = 59 \pm 50°$/s. These values are consistent with those reported in the example of Fig. S3*C* for which the mean angular speed $\langle|\omega|\rangle \simeq 53°$/s before the beginning of the *BS* ($t \leq -0.1$ s) and $\langle|\omega|\rangle \simeq 75°$/s after the end of the reorientation phase ($t \geq 0.4$ s).

As seen in Fig. S3*D*, the *BS* motion starts after the cell has strongly decelerated, 100 ms before its immobilization at $t = 0$. The *BS* phase is characterized by an anti-alignment between the motion unit vector $\hat{\mathbf{m}}$ and the orientation unit vector $\hat{\mathbf{o}}$, highlighted by the pink and purple regions in Fig. S3*B*. After some duration of "pure" *BS*, the cell starts to rotate with an angular speed that increases up to a maximum at time $t = 0.18$ s, that coincides with a minimum of the speed and the end of the *BS*: the cell performs a rotation around its posterior end while being almost immobile (purple regions of Fig. S3). Finally, the rotation slows down while the cell accelerates forward (blue region) until it reaches again its typical speed.

### IV. OUT-OF-PLANE REORIENTATIONS

As mentioned in the *Results* section of the main text, due to the geometry of the quasi-2D environment, the preferred direction of motion of the cells is in the observation plane. However, since the height of the confinement ($h = 340\,\mu$m) is more than twice the length of the cell and ten times its width, the cell is still able to perform 3D motions such as its characteristic helicoidal motion or out-of-plane ($OOP$) reorientation events. Each cell is fitted by an ellipse of time-dependent semi-major axis $a(t)$ and eccentricity $e(t)$. When a cell moves outside of the observation plane, its eccentricity decreases. Therefore, $OOP$ events are detected as time intervals during which $e(t) < e_{thr} = 0.75$, where the threshold $e_{thr}$ is set empirically. Without pillars ($\Phi = 0$), the total duration of $OOP$ events represent $1.2 \pm 0.35\%$ (mean $\pm$ SEM, $T_{obs} = 134\,$min, $M = 10$ experiments) of the total observation time $T_{obs}$. This confirms the observation that typical trajectories are mostly 2D. We also measured that for cells navigating in an environment with pillars ($\Phi \neq 0$), the proportion of those that did reorient outside of the observation plane was not significantly different from the $\Phi = 0$ case. This suggests a negligible influence of the pillars on the 3D motions, which are mostly due to spontaneous reorientations in the bulk. When focusing on $AR$s, the proportion of $AR$ events for which the cell reorients upward or downward instead of left or right in the observation plane, is $19 \pm 3.3\%$, which highlights the three dimensionality of the cells motion during an $AR$. To analyze the $OOP$-$AR$, we first inferred the $OOP$ angle $\theta_\perp(t) = \arccos\sqrt{\frac{a_\parallel^2 - a(t)^2}{a_\parallel^2 - b_\parallel^2}}$ which is the angle between the cell's long axis and the observation plane, where $a_\parallel$ and $b_\parallel$ are respectively the cell's true average semi-major and semi-minor axes. The $BS$ and reorientation phases of the $OOP$-$AR$ were then detected in the same way as for the in-plane $AR$, described in the *Materials and Methods* section of the main text.

### V. DETAILED DYNAMICS OF A SCATTERING EVENT

In the case of a scattering event ($SE$), the contact interaction can be decomposed into 3 successive events: first, a collision with the pillar, then a reorientation and finally a sliding phase, as shown on the typical example of Fig. S4*A*. At the instant of the collision (Fig. S4*B*, $t \sim 40\,$ms), the speed of the cell $v$ is minimum. Follows the reorientation phase (for $t \in [0.04, 0.18]\,$s) during which $v$ increases and $|\omega|$ is non-zero (Fig. S4*C*). The cell then slides against the pillar, at a constant speed $v \simeq 806 \pm 23\ \mu$m/s (Fig. S4*B*, $t \in [0.2, 0.28]\,$s) and with a constant orientation $\theta \simeq -15 \pm 1.0\ ^\circ$/s (Fig. S4*C*, $t \in [0.2, 0.28]\,$s), *i.e.* with a zero angular velocity. Note that its speed during sliding is the same as the one it had prior to the $SE$ away from the pillar (see the first point in Fig. S4*B*). During sliding against the pillar, one could also expect $r_{min}(t)$, the distance between the center of the pillar and the ellipse that fits the cell at time $t$, to be constant since the cell cannot penetrate the pillar.

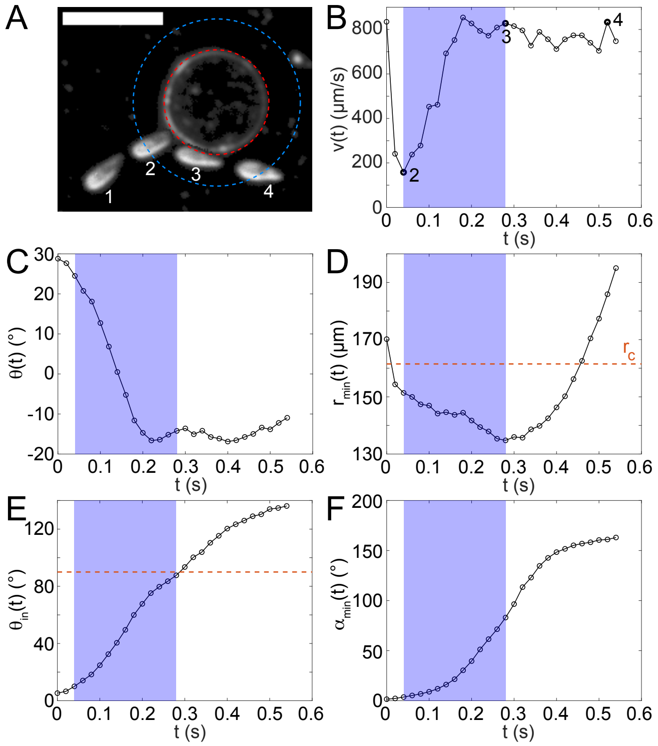


FIG. S4. Detailed dynamics of an example $SE$. *(A)* Composite dark-field image showing a close-up view of a paramecium performing a $SE$ obtained by overlapping images taken at different times separated by 240 ms. Successive positions of the cell are numbered from 1 to 4. At position 2 (*resp.* 3), the cell is at the beginning (*resp.* end) of the contact phase $t_c^i$(*resp.* $t_c^f$). The dashed blue (*resp.* red) circle delimits the interaction (*resp.* contact) corona. The scale bar is $300\,\mu$m long. *(B-F)* Time evolution of the parameters characterizing the trajectory of the cell during the $SE$ of Fig. S4*A*. *(B)* Instantaneous speed of the cell $v(t)$. The origin of time $t = 0$ is taken as the instant $t_{in}$ at which the cell center intercepts the interaction corona. The blue shaded region corresponds to the contact phase $[t_c^i, t_c^f]$. Labels 2 to 4 correspond to the numbered positions of Fig. S4*A*. Note that label 1 is not shown, as it corresponds to $t = -0.2\,$s. *(C)* Orientation $\theta(t)$ of the cell relative to the horizontal axis. *(D)* Distance $r_{min}(t)$ between the center of the pillar and the elliptic fit to the cell. The red dashed line corresponds to the radius of the contact corona $r_c$. *(E)* Incident angle $\theta_{in}(t)$. The red dashed line corresponds to $\theta_{in} = 90^\circ$ at which the contact ends. *(F)* Ellipse angle $\alpha_{min}(t)$ for which $r_{min}$ is reached (see Fig. S2): $\alpha_{min} = 0$ means that the closest point to the pillar center is at the front of the cell. For all graphs, data points are plotted every 20 ms.

Clearly, this is not what is measured (Fig. S4*D*). This can be interpreted as a direct consequence of the left-right asymmetry of the cell combined with its rotation around its long axis, resulting in fluctuations of the elliptic fit. Finally, the cell leaves the pillar when it is tangent to it, *i.e.* when $\theta_{in} = 90^\circ$, which occurs at $t = 0.28\,$s for this example $SE$ (Fig. S4*E* and Fig. S4*A*, position number 3).

### VI. MECHANICAL CONTACT DETECTION

As briefly described in the *Materials and Methods* section of the main text, mechanical contacts between *Paramecium* and a pillar were identified using two criteria that have to be met and the instant of the collision was determined based on a speed measurement. The first criterion is based on a cell-pillar distance measurement, *i.e.* the cell contour has to be within a given contact corona for a contact to be considered. The second criterion is that, when the cell enters the contact corona, its orientation with the local radial direction has to be smaller than a given threshold. Once both criteria are met, the instant of the collision $t_c^i$ between the cell and the pillar is found using a minimum speed measurement. We now detail how both the radius of the contact corona and the orientation threshold were determined.

The radius of the contact corona $r_c$ was obtained by visual inspection of passive interaction events with and without contact. It yielded $r_c = r_p + \delta r$ with $\delta r = 19\,\mu$m, about 5 times the pixel size ($3.81 \pm 0.01\,\mu$m). We checked *a posteriori* the relevance of this empirical value by computing the probability density function (*PDF*) of $r^*_{min} = min\{r_{min}(t)|t \in [t_i, t_o]\}$, which defines the closest distance between the ellipse and the pillar center that is reached within the time interval $[t_i, t_o]$ of a given interaction event. Recall that $t_i$ (*resp.* $t_o$) is the time at which the cell center enters (*resp.* leaves) the interaction corona. For all passive interaction events, the *PDF* of $r^*_{min}$ shows two distinct parts (Fig. S5*A*, black stars): in the interval $[120, \sim 160]\,\mu$m, $r^*_{min}$ is normally distributed (see the solid black line which is a Gaussian fit of the data, with mean value $m_c = 142\,\mu$m and standard deviation $\sigma_c = 7.2\,\mu\text{m}, R^2 = 0.99$), while in the interval $[\sim 160, 220]\,\mu$m, it is uniformly distributed. The first part of the *PDF* of $r^*_{min}$ is a direct signature of the contact interaction between the cell and the pillar. Its mean value $m_c = 142\,\mu$m is close to the independently measured pillar radius $r_p = 142.5\,\mu$m and its standard deviation $\sigma_c = 7.2\,\mu$m is a consequence of the errors in the fitting procedure of the cell contour with an ellipse. The second part of the *PDF* of $r^*_{min}$ is a signature of the interactions without contact and is expected to be uniform. Both parts are separated at $r^*_{min} \approx 160\,\mu$m. This threshold value provides a contact corona width $\delta r \approx 17.5\,\mu$m which is very close to the empirical value we have obtained.

To evaluate the threshold angle below which an interaction event leads to a contact, we evaluated the conditional probability $P(\theta^f_{r_c}|\theta^i_{r_c})$ 2D map, where $\theta^i_{r_c} = \theta_{in}(t^i_{r_c})$ and $\theta^f_{r_c} = \theta_{in}(t^f_{r_c})$ are respectively the incident angle when the cell first enters the corona and when it leaves it, *i.e.* $r_{min}(t) < r_c$ at all times (Fig. S5*B*). One obtains a similar 2D map as the one plotted in Fig. 2*A* of the main text for the interaction corona with two distinct regimes depending on the incident angle $\theta^i_{r_c}$. We evaluated the threshold angle between both regimes by similarly computing the average angle $\langle \theta^f_{r_c} \rangle$ as a func-

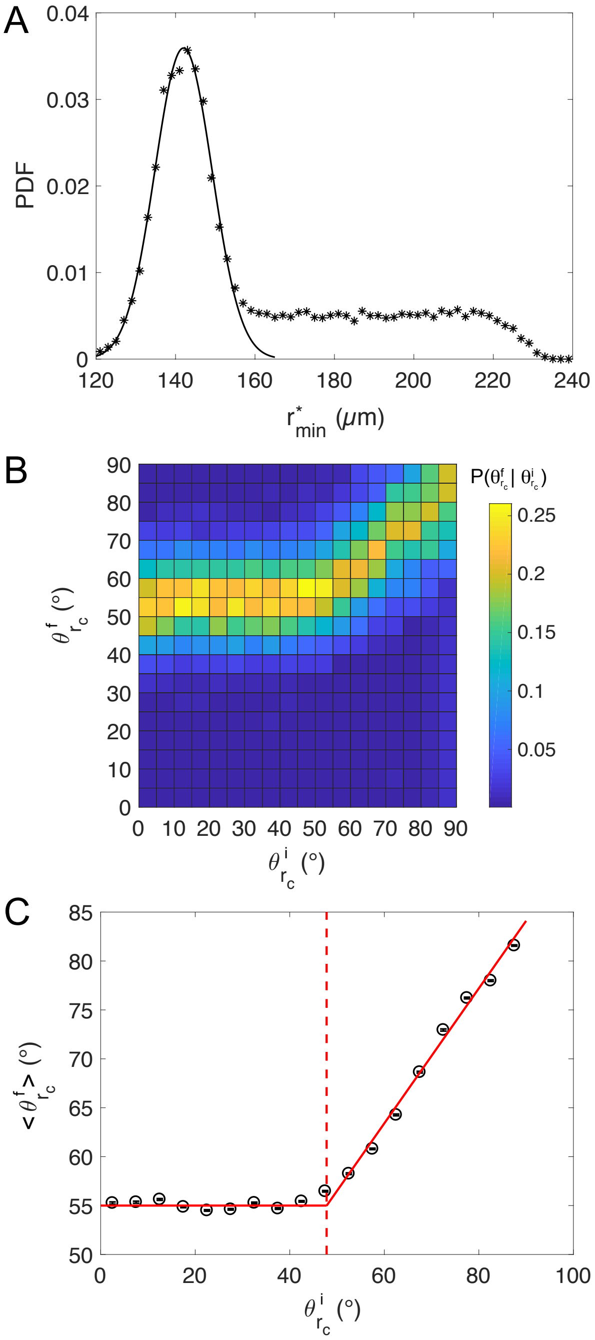


FIG. S5. Mechanical contact detection. *(A)* Probability density function of $r^*_{min}$, the smallest distance to the pillar during an interaction for all passive interaction events (black stars, N = 28031). The bin width is $2\,\mu$m. The solid black line is a Gaussian fit of the *PDF* on the interval $[120; 155]\,\mu$m ($R^2 = 0.99$), yielding a mean value for $r^*_{min}$, $m_c = 142\,\mu$m and standard deviation $\sigma_c = 7.2\,\mu$m. *(B)* 2D plot of the conditional probability $P(\theta^f_{r_c}|\theta^i_{r_c})$ for $\theta^i_{r_c}$ and $\theta^f_{r_c}$ varying between 0° and 90°. Each square bin has a side length of 5°. This plot was obtained for $N = 28031$ passive interaction events. *(C)* Mean output angle $\langle \theta^f_{r_c} \rangle$ versus $\theta^i_{r_c}$, where $\theta^i_{r_c}$ is taken as the center value of each bin. Red solid lines are linear fits of both regimes, intercepting at $\tilde{\theta}^i_{r_c} = 47.8°$, indicated by the vertical red dashed line.

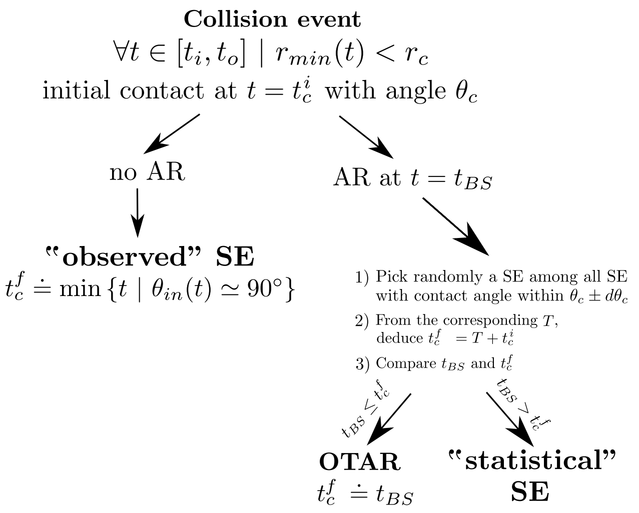


FIG. S6. Sketch of the algorithm used to discriminate *SE*s and *OTAR*s.

tion of $\theta^i_{r_c}$ (Fig. S5*C*, black circles). Like in Fig. 2*B* of the main text, this curve can be modeled and fitted with the following function (solid red line)

$$\left\langle \theta^f_{r_c} \right\rangle (\theta^i_{r_c}) = \begin{cases} \tilde{\theta}^f_{r_c} & \text{if } \theta^i_{r_c} < \tilde{\theta}^i_{r_c} \\ m\theta^i_{r_c} + q & \text{if } \theta^i_{r_c} \geq \tilde{\theta}^i_{r_c} \end{cases}$$

yielding $\tilde{\theta}^f_{r_c} = 55°$ (interval $[0, 40]°$, $R^2 = 0.99$) and $m = 0.69$ and $q = 21.8°$ (interval $[50, 90]°, R^2 = 0.99$). The intersection of both regimes gives the threshold value $\tilde{\theta}^i_{r_c} = 47.8°$ (Fig. S5*C*, vertical red dashed line), below which an interaction event leads to a contact and above which no physical contact between the cell and the pillar exists. We thus consider, in agreement with our visual inspection of a large number of events, that an interaction event leads to contact *if and only if* $\theta^i_{r_c} \leq \tilde{\theta}^i_{r_c}$. Finally, when both criteria discussed above are met, the contact lasts from the first speed minimum on the interval of times where $r_{min} < r_c$, until the cell is tangent to the obstacle.

### VII. DISCRIMINATION OF SCATTERING EVENTS AND OBSTACLE TRIGGERED AVOIDING REACTIONS

Figure S6 sketches the algorithm we have used to discriminate *SE*s and *OTAR*s. As a reminder, a contact interaction starts at $t^i_c$, the collision time, that is defined as the first minimum in the speed of the cell over the time interval during which the cell is inside the contact corona, *i.e* over $[t^i_{r_c}, t^f_{r_c}] \doteq \{t \in [t_i, t_o] \mid r_{min}(t) < r_c\}$. In addition, as explained in the *Materials and Methods* section of the main text and above, the incident orientation angle when entering the contact corona must obey $\theta^i_{r_c} < \tilde{\theta}^i_{r_c}$.

Due to intrinsic limitations of our experiments, the contact could not be resolved visually. While it is not an issue at $t^i_c$, because the transition from the free swimming phase to the contact phase induces a sharp speed decrease and an angular deflection that is easy to detect, no criterion was found to detect the end of the contact $t^f_c$ because the transition from the contact phase to free swimming happens at constant speed and constant orientation as shown in Fig. S5*C-E*. Instead, we decided to postulate that the contact ends when the cell is tangent to the obstacle, *i.e.* when $\theta_{in}(t) \simeq 90°$. However, this reasonable assumption based on geometrical considerations only holds in the case of a cell passively sliding against the pillar. When an *AR* is triggered against the pillar, the cell reorients and leaves the pillar without being necessarily tangent to the pillar.

To solve this problem, we divided the contact interactions into two categories: those without *AR* during $[t^i_{r_c}, t^f_{r_c}]$ (called "observed" *SE*) and those with an *AR*. For the "observed" *SE*, a contact duration is computed directly using the geometrical criterion to define $t^f_c$. On the other hand, for a contact event happening at $t^i_c$ with a collision angle $\theta_c$ and for which an *AR* is triggered at $t_{BS}$, the contact end time is defined as follows.

First, the contact duration $T$ is randomly picked among a pool of $T$ data that is obtained by selecting the "observed" *SE* having a collision angle in the interval $\theta_c \pm d\theta_c$, with $d\theta_c = 5°$. Then, knowing $t^i_c$, $t^f_c = T + t^i_c$ is deduced. Comparing $t_{BS}$ to $t^f_c$ allows to define three cases:

- (i) if $t_{BS} < t^f_c$ and $t_{BS} - t^i_c \in [0, 40]$ ms, it is an "instantaneous" *OTAR*;
- (ii) if $t_{BS} \leq t^f_c$ and $t_{BS} - t^i_c \in ]40, 600]$ ms, it is a "delayed" *OTAR*;
- (iii) if $t_{BS} > t^f_c$, the event is considered to be a "statistical" *SE*.

Note that in the case of an *OTAR*, after classification, the contact end time is redefined as $t^f_c \doteq t_{BS}$, so that the charging of the membrane ends as soon as an *AR* is triggered.

Applying this method over all data ($\Phi \neq 0$, random and square lattices) yielded 15601 *SE*s (with 14729 "observed" *SE*s and 872 "statistical" *SE*s), 670 "instantaneous" *OTAR*s and 1120 "delayed" *OTAR*s. From this, the overall *OTAR* probability was $P(OTAR) = 0.1$.

### VIII. MEASURING THE CONTACT DURATION FOR THE *PAWN* MUTANT SCATTERING EVENTS

As a control, we used *Pawn* mutants, which cannot perform *AR*s due to a lack of voltage-gated calcium channels in the cilia. The mean *AR* frequency measured by our algorithm in the absence of pillars was $\left\langle f^{\phi=0}_{AR} \right\rangle \simeq$

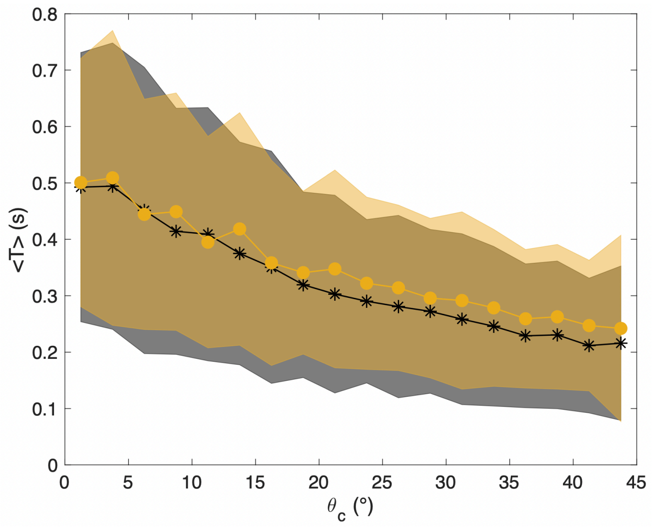


FIG. S7. Average contact duration $\langle T(\theta_c)\rangle$ for the *SE*s of the wild-type strain (black stars) and for the *SE*s of the *Pawn* mutant strain (yellow disks). Data for the *Pawn* strain were compiled from experiments in random environments ($M = 7$ with $\phi = 0.083$, $M = 9$ with $\phi = 0.14$ and $M = 3$ with $\phi = 0.28$) yielding a total of 5274 *SE*s. Note that for the wild-type strain, data are the same as those shown in Fig. 4C of the main text, but with a different bin width. Error bars correspond to the *SEM*.

$4.5\,10^{-3}$ Hz, about 40 times smaller that for the wild-type paramecia, $\left\langle f_{AR}^{\phi=0}\right\rangle \simeq 0.165$ Hz. This shows that our algorithm has a low rate of false positives. With pillars, *Pawn* mutants cannot perform any *OTAR*s nor *AR*s and thus display only *SE*s. In this case, we measured the same trend for the average contact duration $\langle T(\theta_c)\rangle$ (Fig. S7).

### IX. DESCRIPTION OF THE PROVIDED MOVIES

- **Movie S1:** This movie shows a typical spontaneous *AR* in the bulk of a square lattice with $\Phi = 0.14$. The video is played back 2 times slower than real time (the real frame rate is 50 fps).
- **Movie S2:** This movie shows a typical *SE* in a square lattice with $\Phi = 0.14$. The video is played back 2 times slower than real time (the real frame rate is 50 fps).
- **Movie S3:** This movie shows a typical instantaneous *OTAR* in a random lattice with $\Phi = 0.083$. The video is played back 2 times slower than real time (the real frame rate is 50 fps).
- **Movie S4:** This movie shows a typical delayed *OTAR* with $\tau = 0.16$ s in a random lattice with $\Phi = 0.083$. The video is played back 2 times slower than real time (the real frame rate is 50 fps).
